# FASTQuick: Rapid and comprehensive quality assessment of raw sequence reads

**DOI:** 10.1101/2020.06.10.143768

**Authors:** Fan Zhang, Hyun Min Kang

**Author notes:** To whom correspondence should be addressed. **Corresponding Author:** Fan Zhang, Department of Computational Medicine and Bioinformatics, University of Michigan Medical School, 100 Washington Ave, Ann Arbor, MI 48109-2218.

## Abstract

**Background:** Rapid and thorough quality assessment of sequenced genomes in an ultra-high-throughput scale is crucial for successful large-scale genomic studies. Comprehensive quality assessment typically requires full genome alignment, which costs a significant amount of computational resources and turnaround time. Existing tools are either computational expensive due to full alignment or lacking essential quality metrics by skipping read alignment.

**Findings:** We developed a set of rapid and accurate methods to produce comprehensive quality metrics directly from raw sequence reads without full genome alignment. Our methods offer orders of magnitude faster turnaround time than existing full alignment-based methods while providing comprehensive and sophisticated quality metrics, including estimates of genetic ancestry and contamination.

**Conclusions:** By rapidly and comprehensively performing the quality assessment, our tool will help investigators detect potential issues in ultra-high-throughput sequence reads in real-time within a low computational cost, ensuring high-quality downstream analysis and preventing unexpected loss in time, money, and invaluable specimens.

## Findings

### Introduction

Efficient and thorough quality assessment from deeply sequenced genomes in an ultra-high-throughput scale is crucial for successful large-scale sequencing studies. Delay or failure in detecting contamination, sample swaps, quality degradation, or other unexpected problems in the sequencing or library preparation protocol can result in enormous loss of time, money, and invaluable specimens if, for example, hundreds or thousands of samples are found to be contaminated weeks or months later. Ensuring comprehensive quality control of sequence data at real-time speed will assure generation of high-quality sequence reads, and subsequently successful outcomes in the downstream analyses.

Existing quality assessment or quality control (QC) tools mainly fall into two categories – pre-alignment and post-alignment methods – based on whether they require full alignment of the genome prior to the quality assessment. Pre-alignment methods, such as *FASTQC*[1], *PIQA*[2], and *HTQC*[3], produce read-level summary statistics that can be obtained from sequence reads, such as base compositions, k-mer distributions, base qualities, and GC bias levels. However, these pre-alignment methods do not estimate many key quality metrics required for comprehensive quality assessment. These missing metrics include mapping rate, depth distribution, fraction of genome covered, sample contamination, or genetic ancestry information. Other post-alignment methods, such as *QPLOT*[4], *Picard*[5], *GotCloud*[6], and *verifyBamID*[7], provide a subset of these key quality metrics but require full alignment of sequence reads, which typically takes hundreds of CPU hours for deep (e.g.,>30x) sequence genome. (Table 1)

**Table 1.**
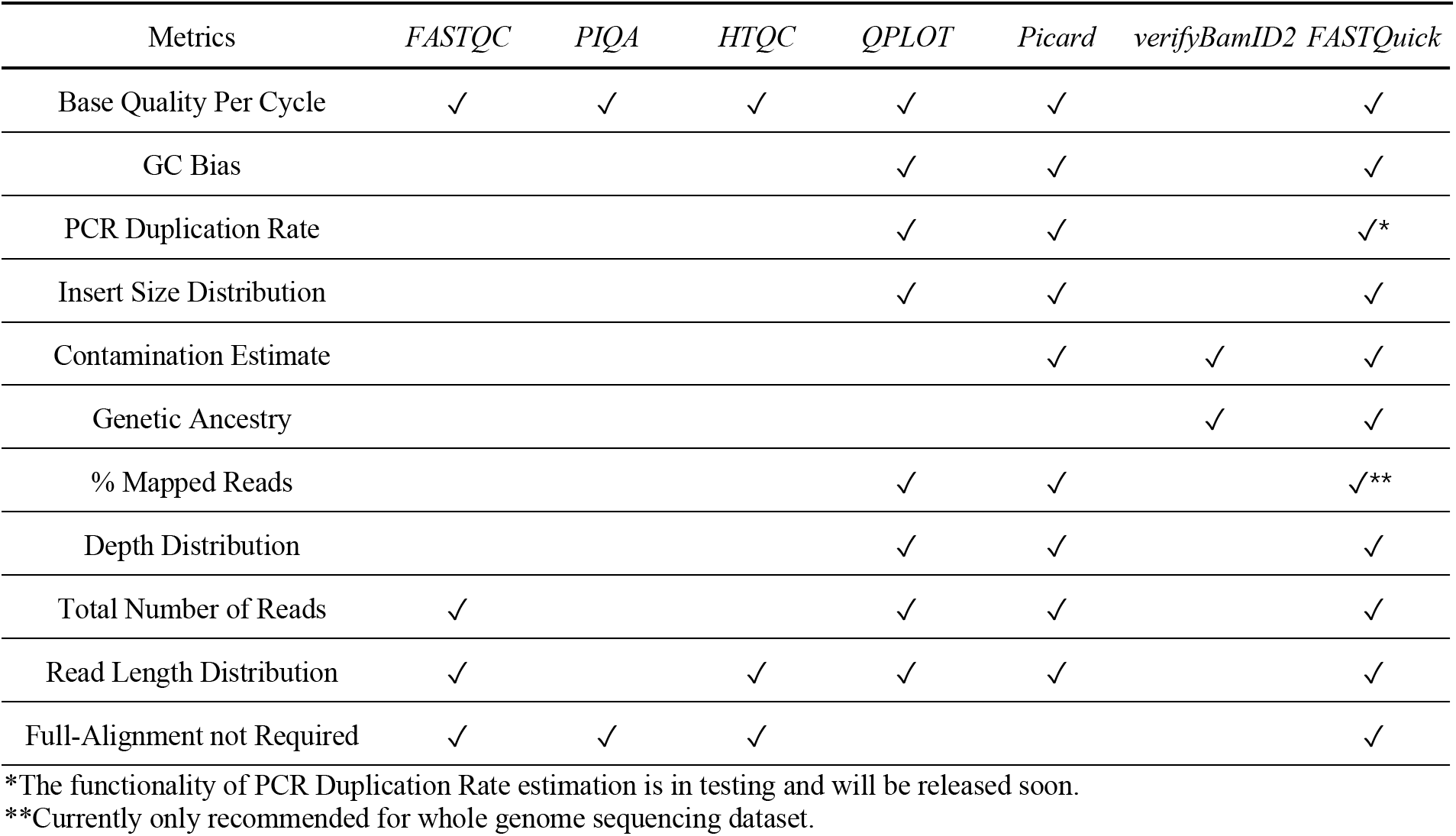
Quality assessment metrics provided by different QC tools

We describe *FASTQuick*, a rapid and accurate set of algorithms and software tools, to combine the merits of QC tools from both categories. By focusing on a variant-centric subset of a reference genome(reduced reference genome), our methods offer up to 30~100-fold faster turnaround time than existing post-alignment methods for deeply sequenced genome, while providing a comprehensive set of quality metrics comparable with *QPLOT* and *verifyBamID* (full-alignment based results from these two tools together constitute most of the important QC metrics from *GotCloud*-based QC pipeline which we will compare against frequently later) with the help of statistical adjustments to account for the reduced reference genome.

### Computational Efficiency

The primary goal of *FASTQuick* is to achieve comprehensive QC with much less computational cost than full-alignment-based QC procedures. A large fraction of the computational gains come from the usage of the reduced reference genome and filtering of unalignable reads through mismatch-tolerant spaced k-mer hashing(Figure 1A)[8]. Compared to alignment to the full human reference genome, aligning a 3x HG00553 genome on the reduced reference genome reduced the run time by 34.9-fold (94,020 vs. 2,697 seconds) using the same algorithm. Using hash table built from mismatch-tolerant spaced k-mers, more than 90% of unalignable reads can be filtered out with very few loss (Table S1) of alignable reads, when 3 or more hits are required (default parameter) for a read to be considered as alignable, saving additional 65% of computational time (Figure 1B). Putting them together, the alignment step of *FASTQuick* (with default parameters) was 100-fold faster (94,020 vs. 939 seconds) than the full genome alignment. We observed that >99% of unalignable reads could be filtered out with a more stringent threshold (7 or more hits) at the expense of 0.01% loss of alignable reads. However, the additional computational gain was only 14% (939 vs. 811 seconds).

**Figure 1.**
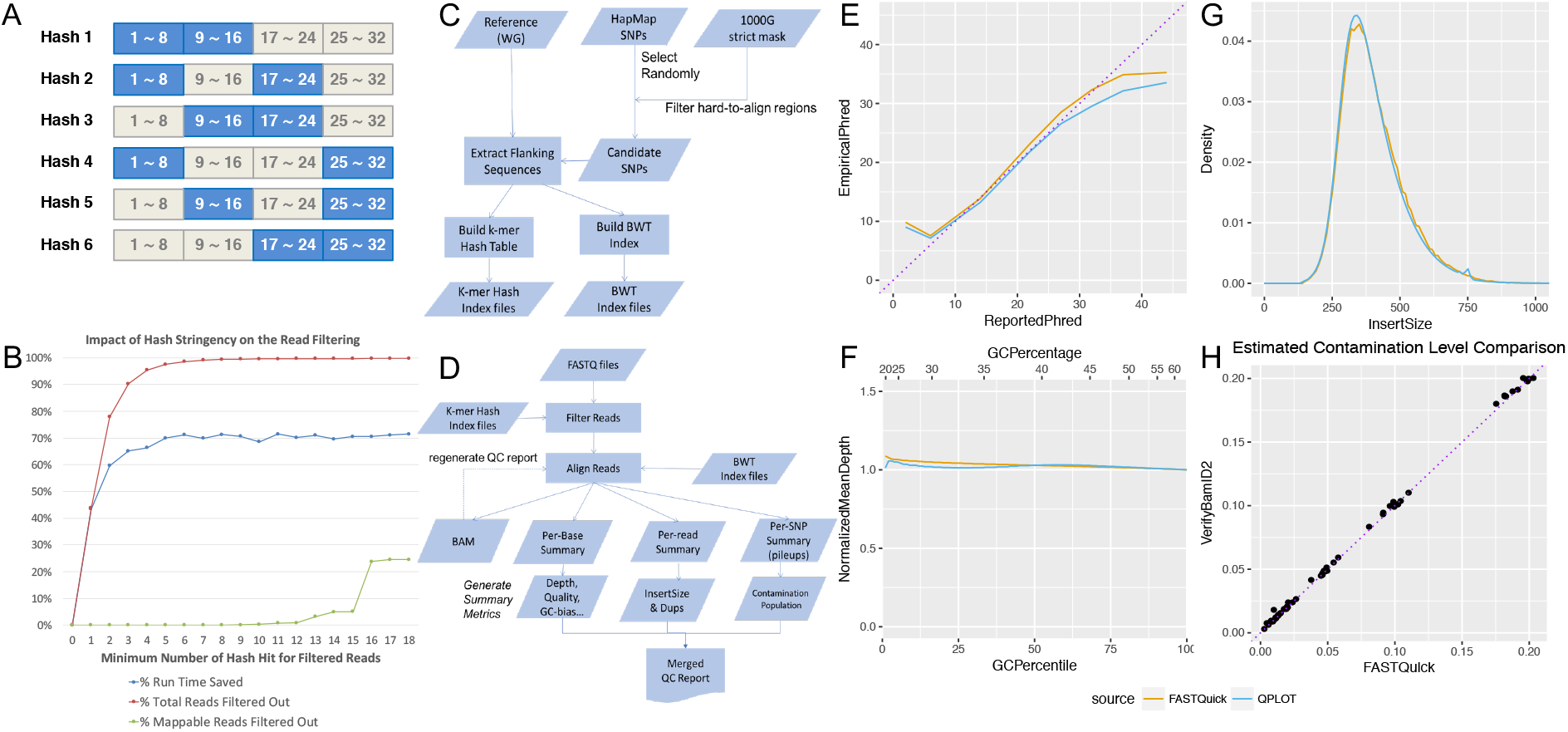
Illustration of *FASTQuick*. **A)** Spaced k-mer hash filter design with the tolerance of mismatches for each 32-mer. **B)** Effect of minimum spaced k-mer hits to be considered for *BWA* alignment on the overall runtime, fraction of total reads filtered, and fraction of falsely filtered alignable reads. k = 3 was used in our experiment. **C)** Procedure to build *FASTQuick* indices with a reduced reference genome for spaced k-mer hash and the *BWA* algorithm. **D)** Procedure to process sequence reads and produce QC metrics using *FASTQuick*. **E)** Comparison of visualizations of reported base qualities (in Phred scale) and empirical base qualities between *QPLOT* and *FASTQuick* for a 38x genome. **F)** Comparison of visualization of GC bias (in normalized mean depth) between *QPLOT* and *FASTQuick* for a 38x genome. **G)** Comparison of estimated insert size distributions between *QPLOT* and *FASTQuick* (after Kaplan-Meier adjustment) for a 38x genome. H) Comparison of estimated contamination rates in an *in-silico* contaminated 1000G samples between *verifyBamlD2* and *QPLOT*. Purple diagonal dot line is y=x.

We also evaluated the overall computational efficiency between *FASTQuick* and the *GotCloud*-based QC pipeline (typical sequence processing pipeline based on full genome alignment as in 1000 genome project and TOPMed project) on the high-coverage genome (38x) and low-coverage (3x) genomes from the 1000 Genomes Project (Table 2). The results demonstrate that *FASTQuick* produces a comparable set of QC metrics to *GotCloud* with a 30~100-fold faster turnaround time.

**Table 2.** Running time comparison (in hours)

| # of Thread | *FASTQuick* Time |  | *GotCloud* QC Time (with *BWA*) |  |
| --- | --- | --- | --- | --- |
|  | HG00553(3X) | NA12878(38X) | HG00553(3X) | NA12878(38X) |
| 1 | 1.03h | 5.48h | 30.95h | 369.56h |
| 2 | 0.53h | 2.46h | 21.53h | 230.85h |
| 4 | 0.33h | 1.76h | 15.83h | 154.91h |
| 8 | 0.24h | 1.75h | 12.74h | 131.85h |

Running time is evaluated as wall-clock elapsed time on a machine with Intel(R) Xeon(R) CPU (X7560 @ 2.27GHz). Reference indexing time is independent of the input sequence dataset and not included. (It takes 3min20s to index human genome under default settings.)

### QC Metrics Produced by *FASTQuick*

*FASTQuick* can automatically generate and visualize the QC metrics listed in Table S2. Briefly, *FASTQuick* generates three types of generic QC summary statistics – per-base, per-read, and per-variant summary statistics. Per-base summary statistics inform mapping rate, depth distribution, GC-bias, and base quality. Per-read summary statistics allow us to estimate insert size distribution adjusted to account for pair-end alignment bias due to the reduced reference genome. Per-variant summary statistics allow us to estimate DNA contamination rate and genetic ancestry. These summary statistics are combined, jointly analyzed, and visualized into an interpretable and user-friendly quality report shown as in Item S1 and Item S2.

### Accuracy of QC Metrics

We compared the distribution of QC metrics generated from *FASTQuick* with those from *GotCloud* on multiple sequenced genomes. The QC metrics shared between *FASTQuick* and *GotCloud* are listed in Table S2. The visualization QC metrics such as base quality recalibration (Figure 1E), normalized mean depth by GC content (Figure 1F), and depth distribution are very close between *FASTQuick* and *GotCloud*. For example, the two-sample Kolmogorov-Smirnov (KS) test statistics, which quantifies the maximum differences between two empirical cumulative distributions of depth was D = 0.040. Similarly, the Wasserstein-1D Distance, which quantifies the average distance between two cumulative distributions of depth, was W= 0.0038. The Wasserstein distance is a widely used metric to evaluate the similarity between two distributions in Generalized Adversary Network[9]. Even though such differences are statistically significant (mainly because of the very large number of observations), it is arguably a small amount difference typically observed between different QC tools on the same sequence data.

One challenge in quality assessment based on the partial alignment of sequence reads to the reduced reference genome is the estimation of insert size distribution. To systematically correct for biased estimation of insert sizes, we statistically integrated the observed insert sizes across all contigs inverse probability weighting based on the Kaplan-Meier curve[10] (See Methods). Applying our correction produces estimated insert size distribution much closer to that from the full alignment (Figure 1G). The KS-test statistic and the Wasserstein-1D distance were D = 0.60 and W = 0.0591 when using 500bp contigs only, but they reduced to D = 0.18 and W = 0.0170 when using both 500bp and 2,000bp contigs when comparing the insert size distributions between *FASTQuick* and *GotCloud*. When adjusting the insert-size distribution using a Kaplan-Meier estimator, they substantially reduced to D = 0.017 and W = 0.0066.

To evaluate the estimation accuracy of contamination rate and genetic ancestry, we prepared artificially contaminated 1000 Genomes samples *in-silico* (see Methods). Then we compare the estimated contamination rate and genetic ancestry from *FASTQuick* with the estimation from the full-alignment QC pipeline-based result. Our results demonstrate that *FASTQuick* can estimate contamination rate (Figure 1H) and genetic ancestry (Table S3) as accurate as the standard method *VerifyBamID2* relying on the full-alignment result.

## Methods

### Overview of *FASTQuick*

*FASTQuick* first constructs a reduced reference genome from a set of flanking sequences surrounding known SNPs and build a BWT index[11] and mismatch tolerant k-mer hash table(Figure 1C). Once the indices are built, *FASTQuick* rapidly filters out unalignable reads whose first 96-bp have less than 3 hits (out of 18 potential hits, among which 6 hits per 32-mer) against the spaced k-mer hash indices, and align filtered sequence reads to the reduced reference genome using the BWT index (Figure 1D). The small fraction of filtered aligned reads will be stored in binary Sequence Alignment/Map format (BAM) [12]. Next, all the summary statistics that are generated from the aligned reads are collected and jointly analyzed to form various QC metrics that are reported in a user-friendly report in HTML (Item S1).

### Construction of Reduced Reference Genome using Flanking Sequences of SNPs

*FASTQuick* constructs a reduced reference genome based on well-alignable flanking sequences around known common SNPs to enrich the reads that are informative both for genetic inference (e.g., contamination and ancestry) and other genomic quality metrics that require reads alignment. Starting from an arbitrary set of known SNPs, *FASTQuick* randomly selects a designated number of SNPs from known common (MAF>5%) SNP set, such as HapMap3[13], while excluding SNPs near hard-to-align regions (e.g., 1000 genome project strict mask region). *FASTQuick* then constructs reduced reference genome using short flanking sequences of the majority of SNPs (e.g., 90%) and long flanking sequences of the remained SNPs.

### Filtering Unalignable Reads with Mismatch-tolerant Hash

Because the reduced reference genome is a small subset of the whole genome sequence, we expect that only a small fraction of reads will be alignable. However, attempting to align all the reads is still computationally expensive. *FASTQuick* builds a hash-based index to rapidly filter out the reads that are unlikely to be aligned to the reduced reference genome. To make the hash robust against sequencing errors, *FASTQuick* builds six locally sensitive hash tables of 16-mers for each 32-mer (Figure 1A), so that 32-mers with 2 or fewer mismatches can still be guaranteed to match to at least one of the hash tables[8].

*FASTQuick* partitions each sequence read into multiple 32-mers and performs hash lookups for each possible 16-mers. For example, for a 100-bp read, eighteen 16-mers (6 per 32-mer) across three 32-mer will be matched to the hash table. For reads longer than 96-bp reads, only the first 96-bp reads are used. *FASTQuick* will decide to filter out a read or not based on whether the number of matching 16-mers is less than a certain threshold k. For example, if k is 3, reads with less than 7 mismatches are guaranteed to pass the filter, and many other reads with more mismatches will pass the filter. If k is 10, reads with less than 3 mismatches are guaranteed to pass the filter. We chose k=3 based on our experiment based on empirical observations (see Findings).The remained reads will then be aligned by the optimized BWA-like algorithms to the reduced reference genome.

### Generating Base-level, Read-level, and Variant-Level QC Metrics

Using the reads aligned to the reduced reference genome, *FASTQuick* generates a full list of base-level, read-level, and variant-level QC metrics (Table S2). Base-level metrics, such as base quality, and sequencing cycle, are recorded directly without using the alignment information. Because the reads spanning the end of flanking sequences may be poorly aligned, *FASTQuick* produces metrics only on the fully alignable portion of flanking sequences. Let the length of the flanking sequence be *w*, and the read length be *r*, then only *2*(w-r) +1* bases spanning the variant site will be considered when calculating base-level summary statistics. Read-level QC metrics, such as the fraction of mapped reads, insert size distribution are estimated and reported based on reads alignment result. Variant-level metrics are collected after alignment result become available and are reported as pile-up bases, estimation of contamination level, and genetic ancestry.

### Bias-Corrected Estimation of Insert Size Distribution

The insert size distribution is typically estimated from distances between the aligned pairs of reads from the fully aligned reads. When using a reduced reference, a large proportion of paired reads may not be fully mapped, and the read pairs that have shorter insert sizes are more likely to be mapped in both ends. As a result, estimating insert size distribution based only on the reads where both ends are mapped will result in biased estimates of insert sizes, as empirically demonstrated using the 38x genome in Figure S1.

We first attempted to resolve this challenge by extending 10% of the variant-centric contigs to be sufficiently long (2000bp), and by estimating insert size only from the reads mapped to longer contigs. This way, we prevent the reduced reference genome from becoming too large to achieve computational efficiency and keep the insert size estimation less biased at the same time. But due to the limited number of long-flanking variants, bias and fluctuations still exist in the estimated insert size distribution. (Figure S2)

To infer insert size distribution more accurately, *FASTQuick* further corrects for the bias nonparametrically using the Kaplan-Meier estimator. Due to the limited length of flanking sequences in the reduced reference, the observed distribution of insert sizes obtained from the reads that both ends are mapped will be biased towards smaller values. To recover the full distribution of insert sizes adjusting for the “censored” reads (i.e., reads with only one of the paired-ends aligned) enriched for large insert sizes, we adopted the Kaplan-Meier estimator as an inverse-probability-of-censoring weighted average[10] as described below.

Specifically, we define a tuple (*t_o_, t_l_, t_r_*) (Figure S3)for each mapped DNA segment (or read pair), where *t_o_* is the observed insert size, *t_l_* is the maximal insert size of *read 1*, and *t_r_* is the maximal insert size of *read 2*. The maximal insert size is defined as the distance between the leftmost/rightmost base of *read 1/read 2* and the rightmost/leftmost base of the flanking region sequence, respectively. This tuple is fully specified only when a read pair is properly aligned, otherwise, for single-end mapped read pair(including partially mapped pair) only one of the two maximal insert sizes (*t_l_* or *t_r_*) is available and unobserved value is set to missing, the rest of the read pairs, such as read pairs that are mapped to different contigs, with low mapping quality, or in abnormal orientation, are discarded in the estimation of insert size distribution. Empirically, given *N* properly aligned read pairs (i.e., tuples without missing values), we can estimate insert size by counting the frequency of different observed insert sizes, *t_o_*, and the cumulative distribution of insert size hence becomes:

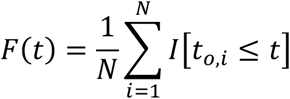

However, as mentioned above, this direct estimation will be severely biased because of reads mapped only in a single end is more likely to have larger insert sizes. To correct for this bias, we use an approach analogous to the estimation of survival function as *S*(*t*) = 1 − *F*(*t*). We can view the leftmost/rightmost base on each flanking region as the start time point, the exact insert size *t_o_* as the time when it fails to observe the data point, and the maximal insert size, *t_l_* and *t_r_*, as the time when the data point is censored. Let the ordered observed time points *t_o_* and censored time points *t_l_* (or *t_r_*) be *τ*. Denote *o_t_* as the number of observed failure cases, i.e., the number of read pairs that have observed insert size less than or equal to *t*, and also denote *c_t_* as the number of censored cases at time *t*, i.e., the number of single-end mapped read pairs have maximal insert size less than or equal to *t*, then let *I* [*τ_J_* ≥ *t*] be indicator function if *j*-th time point larger than certain time *t*(*j*-th insert size larger or equal to *t*). Then the risk set is:

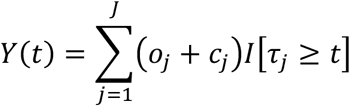

Then the Kaplan-Meier estimator 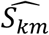 of *S*(*t*):

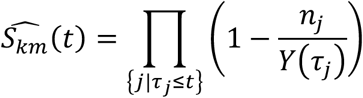

Satten *et al*. [10] proposed a simplified algorithm to iteratively estimate survival function for failure times and survival functions for censoring times, by which we conveniently estimate *F*(*t*).

### Estimation of Contamination Rates and Genetic Ancestry

We also implemented the likelihood-based methods to estimate genetic ancestry and contamination rate in *FASTQuick*. The details of these methods will be fully described in *VerifyBamID2*[14]. In *FASTQuick*, to seamlessly integrated these methods into our ultra-fast QC procedure, we designed compatible variant-centric data structures and input/output interfaces that can directly deliver sequence information and estimated statistics from *FASTQuick* to modules that estimate contamination and genetic ancestry.

### Support for Target Sequencing Dataset

*FASTQuick* also has provided options to incorporate target regions. We can conveniently use the exome region list for Exome-seq, and abundantly expressed gene list for RNA-seq as input information to only select markers within the list. We prepared the result generated by *FASTQuick* for exome sequencing data of HG00553 from the 1000 genome project as a demonstration (Item S2).

## Discussion

We described *FASTQuick*, which addresses computational challenges in quality control of ultra-high-throughput sequence data, by focusing on sequence reads mappable to an informative subset of the reference genome. Our results demonstrate that *FASTQuick* achieves with on average 30 ~ 100-fold faster turnaround time than methods based on full sequence alignment while producing comprehensive and accurate QC metrics. Compared to previous quality assessment methods that do not align sequence reads at all, *FASTQuick* provides more comprehensive QC metrics such as depth distribution, insert size distribution, contamination, and genetic ancestry.

*FASTQuick* leverages several methods, such as spaced-kmer hash table and Kaplan-Meier estimator, to enable rapid and accurate estimation of QC metrics. Interestingly, the computational time is much faster than the time required to convert and compress Illumina’s BCL formatted files into FASTQ files. Therefore, *FASTQuick* can work as a UNIX pipe during the conversion procedures to increase efficiency in the sequencing pipeline.

There are potential drawbacks of only using the reduced (subset of) reference genome, but *FASTQuick* applies heuristics to avoid such drawbacks. For example, reads that originate from multiple homologous regions on the genome may be misaligned to the same contig on the reduced genome, which may affect variant-level quality metrics. *FASTQuick* addresses this issue by strictly selecting regions that are unique and easy to align (callable regions), and we demonstrated the effectiveness by showing that contamination and genetic ancestry estimates are almost identical to the estimation from full genome alignment result. Another issue could be the excessive single-end alignment, for example, it will skew the estimation of insert size distribution toward smaller value. We applied Kaplan-Meier estimator to correct the estimation as described above. There are still limitations associated with the reduced reference genome. For example, a precise estimation of % mapped reads is challenging, especially for targeted sequencing reads, due to the lack of repetitive sequences. Analysis involving structural variation or comprehensive screening of GWAS variants may not be feasible under *FASTQuick*’s settings.

Currently, *FASTQuick* is only suitable for short sequence reads. To enable an analysis of long sequence reads, additional alignment algorithms such as *Minimap2* [15] could be incorporated. Extending *FASTQuick* to other types of sequence data, such as RNA-seq, ChIP-seq, and ATAC-seq, should also be possible if the technology-specific characteristics are properly considered and accounted for. What’s more, *FASTQuick* can serve as a general down-sampling step prior to analysis like sample-swap detection, kinship estimation with the help of alignment result on common variants. More broadly, although we demonstrated *FASTQuick*’s capability by using human genome analysis as an example, the whole pipeline is adaptable easily to other organisms provided with corresponding genomic databases.

Unlike hardware-accelerated solutions achieve fast speed by introducing specialized hardware, such as *DRAGEN*[16] and *Parabricks*[17], *FASTQuick* gains its speed from optimized algorithms that are specially designed for the reduced genome setting. Compared to omnipurpose proprietary tools like *DRAGEN* and *Parabricks, FASTQuick* is an open-source tool that does not require specific hardware such as GPU or FPGA devices and is specifically designed for quality assessment which can be critical to have rapid turnaround time in sequence analysis workflow and add a great value to the existing sequence analysis ecosystem.

## Supporting information

Supplemental Items

## Availability and requirements

**Project name:** <u>FASTQuick</u>

**Project home page:** https://github.com/Griffan/FASTQuick

**Operating system(s):** <u>Linux, MacOS</u>

**Programming language:** <u>C++, Shell, R</u>

**Other requirements:** <u>CMAKE, libhts, ggplot2, knitr</u>

**License:** <u>MIT</u>

**Any restrictions to use by non-academics:** <u>None</u>

## Declarations

### Ethics approval and consent to participate

Not applicable

### Consent for publication

All the authors consent to publish.

### Availability of data and materials

Datasets are publicly available at the Trans-Omics Precision Medicine (TOPMed) project and the 1000 genome project.

### Authors’ contributions

F.Z contributed to the coding material and experiments. F.Z and H.M.K. together contributed to the writing material.

## Supplementary Materials

### Experimental Data

We selected a deeply sequenced genome of a publicly available sample (NA12878) from the Trans-Omics Precision Medicine (TOPMed) project for most evaluations. Also, we selected an exome-sequencing dataset of the same sample (NA12878) from the 1000 genome project (SRR098401) for target sequencing evaluation. To evaluate computational efficiency for the low-pass sequence genome, we also evaluated another sample (HG00553) from the 1000 Genomes Project (ERR013170, ERR015764, and ERR018525). To evaluate the accuracy of contamination estimates we constructed 10 genomes with *in-silico* contamination by randomly sampling aligned sequence reads from samples in 1000 Genomes phase 3 project and then mixing reads from different samples proportional to the intended contamination rates α ∈ {0.01,0.02,0.05, 0.1,0.2}, as described in *VerifyBamID2*[14].

### Supplementary Tables

**Table S1.** Impact of Mismatch Threshold on Kmer-Hash based Reads Filtering

| Mismatch Threshold K | Total Reads | Filtered Reads | Remained Mappable Reads |
| --- | --- | --- | --- |
| 0 | 96240521 | 0 (0.0000%) | 270405 (100.0000%) |
| 1 | 96240521 | 42200008 (43.8485%) | 270405 (100.0000%) |
| 2 | 96240521 | 74965846 (77.8943%) | 270405 (100.0000%) |
| 3 | 96240521 | 86900055 (90.2947%) | 270405 (100.0000%) |
| 4 | 96240521 | 91827374 (95.4145%) | 270403 (99.9993%) |
| 5 | 96240521 | 93920755 (97.5896%) | 270397 (99.9970%) |
| 6 | 96240521 | 94865507 (98.5713%) | 270381 (99.9911%) |
| 7 | 96240521 | 95392311 (99.1187%) | 270371 (99.9874%) |
| 8 | 96240521 | 95635568 (99.3714%) | 270197 (99.9231%) |
| 9 | 96240521 | 95756594 (99.4972%) | 270105 (99.8891%) |
| 10 | 96240521 | 95828932 (99.5723%) | 269643 (99.7182%) |
| 11 | 96240521 | 95872139 (99.6172%) | 268304 (99.2230%) |
| 12 | 96240521 | 95898966 (99.6451%) | 268160 (99.1698%) |
| 13 | 96240521 | 95941830 (99.6896%) | 261835 (96.8307%) |
| 14 | 96240521 | 95961621 (99.7102%) | 257046 (95.0596%) |
| 15 | 96240521 | 95969319 (99.7182%) | 256664 (94.9184%) |
| 16 | 96240521 | 96026258 (99.7774%) | 206014 (76.1872%) |
| 17 | 96240521 | 96031175 (99.7825%) | 204101 (75.4797%) |
| 18 | 96240521 | 96033063 (99.7844%) | 203998 (75.4417%) |

Mismatch threshold (number of kmer hits) experiments to evaluate the kmer-hash based reads filtering effectiveness

**Table S2.** Summary statistics and visualization items produced by *FASTQuick*

| Output File Name | Visualization | Description |
| --- | --- | --- |
| [output_prefix].AdjustedInsertSizeDist | Y | Adjusted Insert Size Distribution |
| [output_prefix].DepthDist | Y | Depth distribution |
| [output_prefix].EmpCycleDist | Y | Empirical Base Quality vs. Sequencing Cycle |
| [output_prefix].EmpRepDist | Y | Empirical Base Quality vs. Reported Base Quality |
| [output_prefix].GCDist | Y | GC Content Distribution |
| [output_prefix].InsertSizeTable | N | Insert Size for Each Read Pair |
| [output_prefix].Likelihood | N | Genotype Likelihood |
| [output_prefix].Pileup | N | Pileup format information |
| [output_prefix].RawInsertSizeDist | Y | Insert Size Distribution (Unadjusted) |
| [output_prefix].bam | N | Reads Alignment |
| [output_prefix].Summary | N | General Summary Report |
| [output_prefix].pdf | Y | Visualization file containing various QC metrics |
| FinalReport.html | Y | Integrated report including statistics listed above |

**Table S3.** Comparison of Genetic Ancestry Estimation between *FASTQuick* and *VerifyBamID2*.

| Simulated_Sample | Tool | PC1_mu | PC1_sd | PC2_mu | PC2_sd |
| --- | --- | --- | --- | --- | --- |
| HG00097_HG00464 | *FASTQuick* | -0.0102 | 0.0004 | -0.0235 | 0.0029 |
| HG00097_HG00464 | *VerifyBamID2* | -0.0104 | 0.0003 | -0.0234 | 0.0033 |
| HG00097_NA19204 | *FASTQuick* | -0.0100 | 0.0001 | -0.0251 | 0.0001 |
| HG00097_NA19204 | *VerifyBamID2* | -0.0101 | 0.0001 | -0.0250 | 0.0002 |
| HG00105_HG00463 | *FASTQuick* | -0.0101 | 0.0002 | -0.0259 | 0.0017 |
| HG00105_HG00463 | *VerifyBamID2* | -0.0104 | 0.0004 | -0.0247 | 0.0027 |
| HG00105_NA19152 | *FASTQuick* | -0.0096 | 0.0004 | -0.0265 | 0.0003 |
| HG00105_NA19152 | *VerifyBamID2* | -0.0097 | 0.0003 | -0.0267 | 0.0004 |
| HG00692_HG00107 | *FASTQuick* | -0.0167 | 0.0004 | 0.0299 | 0.0023 |
| HG00692_HG00107 | *VerifyBamID2* | -0.0169 | 0.0005 | 0.0306 | 0.0022 |
| HG00692_NA19204 | *FASTQuick* | -0.0165 | 0.0005 | 0.0306 | 0.0014 |
| HG00692_NA19204 | *VerifyBamID2* | -0.0166 | 0.0004 | 0.0306 | 0.0014 |
| HG00708_HG00101 | *FASTQuick* | -0.0165 | 0.0002 | 0.0312 | 0.0005 |
| HG00708_HG00101 | *VerifyBamID2* | -0.0165 | 0.0002 | 0.0312 | 0.0004 |
| HG00708_NA19152 | *FASTQuick* | -0.0161 | 0.0004 | 0.0314 | 0.0003 |
| HG00708_NA19152 | *VerifyBamID2* | -0.0166 | 0.0000 | 0.0312 | 0.0002 |
| NA19141_HG00107 | *FASTQuick* | 0.0355 | 0.0005 | 0.0042 | 0.0002 |
| NA19141_HG00107 | *VerifyBamID2* | 0.0354 | 0.0005 | 0.0042 | 0.0002 |
| NA19141_HG00464 | *FASTQuick* | 0.0353 | 0.0005 | 0.0038 | 0.0008 |
| NA19141_HG00464 | *VerifyBamID2* | 0.0355 | 0.0004 | 0.0039 | 0.0009 |
| NA19190_HG00101 | *FASTQuick* | 0.0338 | 0.0000 | 0.0038 | 0.0002 |
| NA19190_HG00101 | *VerifyBamID2* | 0.0339 | 0.0004 | 0.0042 | 0.0007 |
| NA19190_HG00463 | *FASTQuick* | 0.0341 | 0.0004 | 0.0045 | 0.0005 |
| NA19190_HG00463 | *VerifyBamID2* | 0.0340 | 0.0002 | 0.0040 | 0.0002 |

Comparison of genetic ancestry estimation (via principal component coordinates) between *FASTQuick* and *VerifyBamID2* using 60(12 pairs of datasets with 5 different mixing rate 0.01, 0.02, 0.05, 0.1, 0.2) simulated low coverage sequencing samples from 1000 genome project. *FASTQuick* or *VerifyBamID2* independently estimates the set of PC coordinates of each simulated sample.

### Supplementary Figures

**Figure S1.**
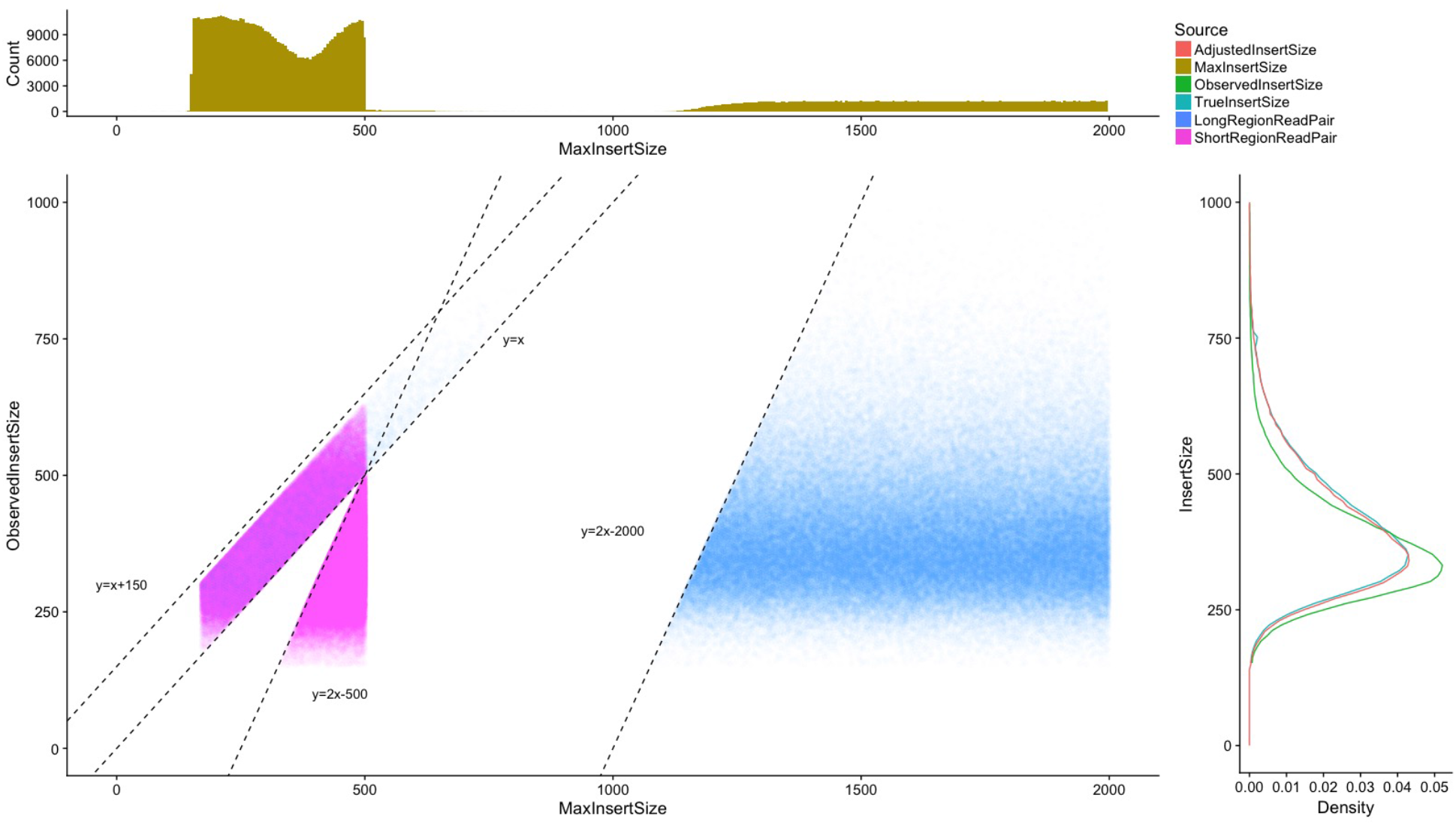
Marginal distribution of max insert size and observed insert size in the reduced genome under 250bp(short) and 1000bp(long) flanking length configuration. Top) Marginal distribution of max insert size. Right) Marginal distribution of observed insert size(green), along with true insert size distribution (Blue) and adjusted insert size distribution(red) Bottom) Scatter plot of read pairs with max insert size and observed insert size being coordinates. Blue dots represent read pairs mapped to the long flanking region; purple dots represent read pairs mapped to the short flanking region. The band between the line “y=x” and line “y=x+150” are read pairs partially mapped. The line “y=2x-500” and line “y=2-2000” are the effective boundaries where read pairs have both ObservedInsertSize and MaxInsertSize for 250bp flanking region and 1000bp flanking region, respectively.

**Figure S2.**
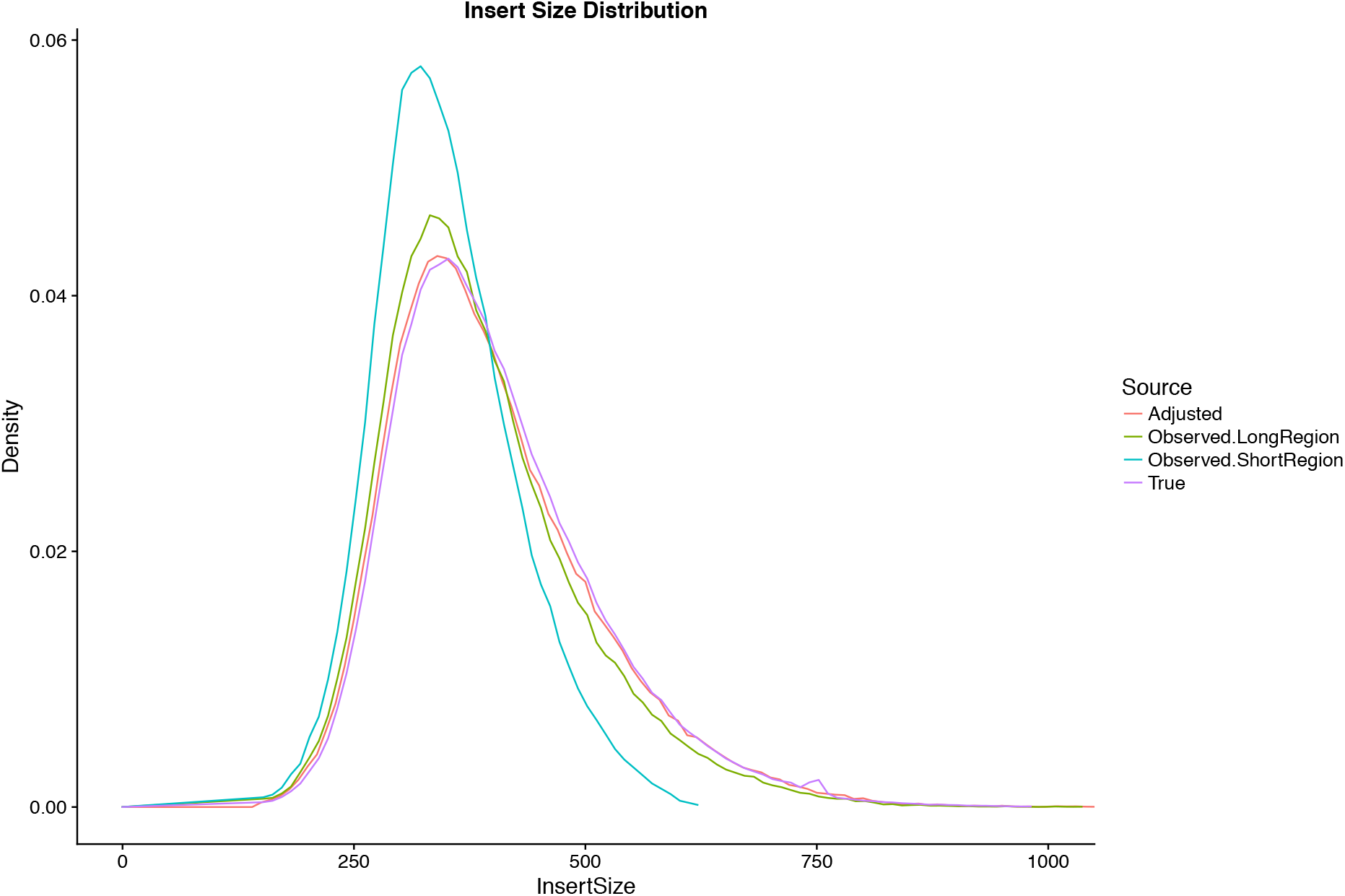
Biased insert size distribution in reduced genome under 250bp(short) or 1000bp(long) flanking length configuration. Each color represents one scenario of insert size estimation without correction. “Observed.LongRegion” (green) is when insert size distribution estimated only using reads mapped to the long flanking region; “Observed.ShortRegion”(blue) is when only using reads mapped to the short flanking region; “True” (purple) is insert size distribution estimated under full genome alignment; “Adjusted”(red) is insert size distribution estimated by *FASTQuick*.

**Figure S3.**
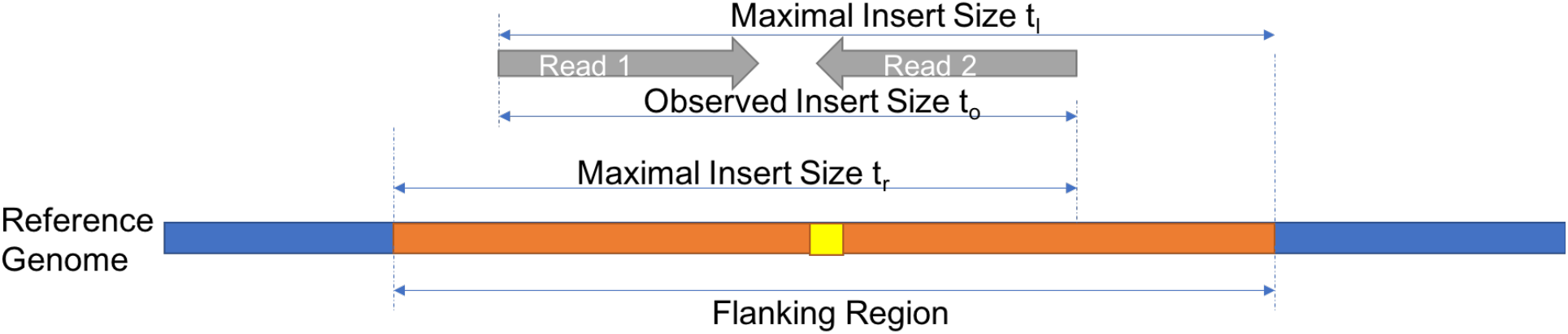
Definition of Insert Size Tuple. The blue portion represents a reference genome backbone. The orange portion represents the extracted flanking region. The yellow portion represents a variant. The gray bars represent a pair of reads aligning to this flanking region.

**Item S1. Detailed Quality Assessment Final Report of HG00553 Whole Genome Dataset (in separate supplementary materials).**

**Item S2. Detailed Quality Assessment Final Report of HG00553 Exome Dataset (in separate supplementary materials).**

