## Supplemental Items for "FASTQuick: Rapid and comprehensive quality assessment of raw sequence reads": Item_S1_FASTQuick_HG00553_low_coverage.FinalReport.html

FASTQuick Summary Report


### FASTQuick Summary Report

- FASTQ File List
- Data Production by FASTQ file
- Depth Distribution
- Summary Statistics
- Summary Plot
- Genetic Ancestry Plot

#### FASTQ File List

FASTQ List Table

| FileIndex | PairEnd1 | PairEnd2 |
| --- | --- | --- |
| 1 | ERR013170\_1.fastq.gz | ERR013170\_2.fastq.gz |
| 2 | ERR015764\_1.fastq.gz | ERR015764\_2.fastq.gz |
| 3 | ERR018525\_1.fastq.gz | ERR018525\_2.fastq.gz |

#### Data Production by FASTQ file

Data Production Table

| FileIndex | NumOfBases | NumOfReads | NumOfUmappedReads | NumOfLowMAPQReads | NumOfQCPassReads | ReadLength |
| --- | --- | --- | --- | --- | --- | --- |
| 1 | 5394785544 | 49951718 | 2342884 | 51223 | 100881 | 108 |
| 2 | 1531077120 | 14176640 | 664757 | 14770 | 28278 | 108 |
| 3 | 3555557208 | 32921826 | 1703961 | 26901 | 60311 | 108 |
| Total | 10481419872 | 97050184 | 4711602 | 92894 | 189470 | 108 |

#### Depth Distribution

#### Summary Statistics

Summary Statistics

| Statistics | Value |
| --- | --- |
| Estimated Read Mapping Rate | 0.896072 |
| Estimated Read PCR Duplication Rate | 0.0529013 |
| Whole Genome Coverage | 3.34074[10481419872/3137454505] |
| Expected Read Depth | 3.61386[10481419872/2900338372] |
| Estimated Read Depth | 3.24109[14931803/4607026] |
| Estimated Percentage of Accessible Genome Covered | 94.3358% |
| Total Accessible Genome Size | 4607026 |
| Depth 1 or above position fraction | 0.943358 |
| Depth 2 or above position fraction | 0.800941 |
| Depth 5 or above position fraction | 0.24105 |
| Depth 10 or above position fraction | 0.00576945 |
| Q20 Base Fraction | 0.94961 |
| Q30 Base Fraction | 0.750469 |
| Estimated AvgDepth for Q20 bases | 3.26257 |
| Estimated AvgDepth for Q30 bases | 2.57838 |
| Median Insert Size(>=500bp) | 518 |
| Median Insert Size(>=300bp) | 456 |
| Contamination Level | 0.00462313 |

#### Summary Plot

```
## Warning: Removed 1 rows containing missing values (geom_path).
```

#### Genetic Ancestry Plot
