## Supplemental Items for "FASTQuick: Rapid and comprehensive quality assessment of raw sequence reads": Item_S2_FASTQuick_HG00553_exome.FinalReport.html

FASTQuick Summary Report


### FASTQuick Summary Report

- FASTQ File List
- Data Production by FASTQ file
- Depth Distribution
- Summary Statistics
- Summary Plot
- Genetic Ancestry Plot

#### FASTQ File List

FASTQ List Table

| FileIndex | PairEnd1 | PairEnd2 |
| --- | --- | --- |
| 1 | SRR070481\_1.fastq.gz | SRR070481\_2.fastq.gz |
| 2 | SRR070780\_1.fastq.gz | SRR070780\_2.fastq.gz |

#### Data Production by FASTQ file

Data Production Table

| FileIndex | NumOfBases | NumOfReads | NumOfUmappedReads | NumOfLowMAPQReads | NumOfQCPassReads | ReadLength |
| --- | --- | --- | --- | --- | --- | --- |
| 1 | 5496822400 | 54968224 | 1083443 | 381075 | 2447313 | 100 |
| 2 | 5462980800 | 54629808 | 1104009 | 383348 | 2434984 | 100 |
| Total | 10959803200 | 109598032 | 2187452 | 764423 | 4882297 | 100 |

#### Depth Distribution

```
## Warning: Removed 1 rows containing missing values (geom_path).
```

#### Summary Statistics

Summary Statistics

| Statistics | Value |
| --- | --- |
| Estimated Read Mapping Rate | 0.463495 |
| Estimated Read PCR Duplication Rate | 0.0966153 |
| Whole Genome Coverage | 3.49322[10959803200/3137454505] |
| Expected Read Depth | 234.754[10959803200/46686362] |
| Estimated Read Depth | 109.338[282211947/2581108] |
| Estimated Percentage of Accessible Genome Covered | 99.2163% |
| Total Accessible Genome Size | 2581108 |
| Depth 1 or above position fraction | 0.992163 |
| Depth 2 or above position fraction | 0.987905 |
| Depth 5 or above position fraction | 0.974027 |
| Depth 10 or above position fraction | 0.94913 |
| Q20 Base Fraction | 0.971707 |
| Q30 Base Fraction | 0.883132 |
| Estimated AvgDepth for Q20 bases | 107.083 |
| Estimated AvgDepth for Q30 bases | 97.3223 |
| Median Insert Size(>=500bp) | 530 |
| Median Insert Size(>=300bp) | 340 |
| Contamination Level | 0.000114851 |

#### Summary Plot

```
## Warning: Removed 1 rows containing missing values (geom_path).
```

#### Genetic Ancestry Plot
